# MGDrivE 3: A decoupled vector-human framework for epidemiological simulation of mosquito genetic control tools and their surveillance

**DOI:** 10.1101/2023.09.09.556958

**Authors:** Agastya Mondal, Héctor M. Sánchez C., John M. Marshall

## Abstract

Novel mosquito genetic control tools, such as CRISPR-based gene drives, hold great promise in reducing the global burden of vector-borne diseases. As these technologies advance through the research and development pipeline, there is a growing need for modeling frameworks incorporating increasing levels of entomological and epidemiological detail in order to address questions regarding logistics and biosafety. Epidemiological predictions are becoming increasingly relevant to the development of target product profiles and the design of field trials and interventions, while entomological surveillance is becoming increasingly important to regulation and biosafety. We present MGDrivE 3 (Mosquito Gene Drive Explorer 3), a new version of a previously-developed framework, MGDrivE 2, that investigates the spatial population dynamics of mosquito genetic control systems and their epidemiological implications. The new framework incorporates three major developments: i) a decoupled sampling algorithm allowing the vector portion of the MGDrivE framework to be paired with a more detailed epidemiological framework, ii) a version of the Imperial College London malaria transmission model, which incorporates age structure, various forms of immunity, and human and vector interventions, and iii) a surveillance module that tracks mosquitoes captured by traps throughout the simulation. Example MGDrivE 3 simulations are presented demonstrating the application of the framework to a CRISPR-based homing gene drive linked to dual disease-refractory genes and their potential to interrupt local malaria transmission. Simulations are also presented demonstrating surveillance of such a system by a network of mosquito traps. MGDrivE 3 is freely available as an open-source R package on CRAN (https://cran.r-project.org/package=MGDrivE2) (version 2.1.0), and extensive examples and vignettes are provided. We intend the software to aid in understanding of human health impacts and biosafety of mosquito genetic control tools, and continue to iterate per feedback from the genetic control community.

**Author summary:** Vector-borne diseases such as malaria cause massive morbidity and mortality throughout much of the world. Currently-available control measures, such as insecticide-based tools and antimalarial drugs, have limited impact and are waning in effectiveness, hence there is a need for novel tools to complement existing ones.

Mosquito genetic control tools, such as gene drive systems and genetic versions of the sterile insect technique, offer a range of promising options, the development of which has greatly expanded since the advent of CRISPR-based gene-editing. Recently, we proposed MGDrivE 2 (Mosquito Gene Drive Explorer 2), which incorporates epidemiology into simulations of the dynamics of these systems in spatially-structured mosquito populations; however, that framework relied on simple model representations of vector-borne diseases. Here, we present MGDrivE 3, which decouples the vector portion of the model from the human portion, allowing the mosquito genetic control framework to be paired with more-detailed epidemiological frameworks. As an example, we implement the human transmission dynamics of the Imperial College London malaria model. We also incorporate a network of mosquito traps for surveillance. As genetic control technology edges closer towards field implementation, more detailed predictions of its epidemiological and biosafety implications are needed. We propose MGDrivE 3 to fulfill this role.

## Introduction

Since the advent of CRISPR-based gene-editing, mosquito genetic control technology has been advancing at a rapid pace, with a plethora of novel genetic constructs being developed in the lab and the prospect of field releases being discussed in earnest. For malaria vectors, recent constructs include a suppression gene drive targeting the *doublesex* gene in *Anopheles gambiae* [1], a replacement gene drive linked to dual antimalarial effector genes in both *An. gambiae* and *Anopheles coluzzii* [2], and a genetic version of the sterile insect technique engineered in *An. gambiae* [3]. As the prospect of environmental releases of constructs like these nears, there is a need for increasingly detailed mathematical models to predict the spread of genes through populations, as well as their epidemiological and biosafety implications [4].

Disease transmission is a key area requiring further development in mosquito genetic control models. Models thus far have tended to emphasize entomological properties and outcomes, such as changes in allele frequencies over time and geographical spread [5–8], and while epidemiological dynamics have sometimes been incorporated, models have tended to utilize simple representations of vector-borne disease transmission, such as the Ross-Macdonald model of malaria transmission, with some exceptions [9]. Meanwhile, detailed models of malaria transmission have been developed by several groups [10–12] incorporating symptomatic and asymptomatic infection, variable parasite density in humans, age structure, mosquito biting heterogeneity, and interventions such as vector control utilizing long-lasting insecticide-treated nets (LLINs) and indoor residual spraying with insecticides (IRS), and antimalarial drug therapy. Incorporating this level of epidemiological detail into mosquito genetic control models would be of great utility considering that genetic control tools will likely be implemented alongside other interventions, expected epidemiological impacts should be a focus in developing these products [4], and initial field trials are expected to have a measured entomological outcome alongside a modeled epidemiological one [13].

Surveillance is another key area requiring inclusion in mosquito genetic control models. Models thus far have tended to record allele frequencies and population densities directly from model output, while incorporating traps explicitly within models would allow questions related to the optimal density and placement of traps to be explored. This would be useful to assess monitoring requirements to both: i) accurately measure effectiveness of genetic control (e.g., establishment and persistence of alleles at future field sites), and ii) detect unintended spread of transgenes beyond the testing or trial site [14]. This latter concern is of particular importance for non-localized gene drive mosquito projects, which have potential to spread on a wide, potentially regional, scale. Efficient, model-informed surveillance programs are therefore essential, as surveillance is expected to be a major cost driver for this technology.

Previously, our group developed MGDrivE (Mosquito Gene Drive Explorer) [5] to model the spatial population dynamics of a variety of mosquito genetic control systems, and MGDrivE 2 [15], incorporating simple models of malaria and arbovirus transmission, seasonality in mosquito populations, and a novel formulation of mosquito and human state space utilizing stochastic Petri nets (SPNs). Here, we present MGDrivE 3, a new version of MGDrivE 2 that incorporates three major developments: i) a decoupled sampling algorithm allowing the vector and human portions of the model to be readily modularized, and hence for the mosquito portion of MGDrivE to be paired with a more-detailed epidemiological framework, ii) a version of the Imperial College London (ICL) malaria transmission model [10, 11], which incorporates age structure, various forms of immunity, human and vector interventions, and more meaningful disease outcomes, and iii) surveillance functionality that tracks mosquitoes captured by traps throughout the simulation. As such, parasite transmission can now be modeled according to mosquito genotype, genetic control interventions can now be modeled alongside other interventions (such as LLINs, IRS and antimalarial drugs), and the dynamical and surveillance implications of mosquito traps can now be modeled.

In this paper, we describe the new features implemented in MGDrivE 3. Additionally, we present an example applying the framework to a hypothetical release of a CRISPR-based homing gene drive system linked to dual disease-refractory genes and their implications for malaria transmission in a low-transmission island setting.

Simulations are also presented demonstrating surveillance of a similar drive system by a network of mosquito traps. We conclude with a discussion of future directions and applications of MGDrivE 3 to the development and application of mosquito genetic control tools towards the goal of vector-borne disease control.

### Design and implementation

As with the MGDrivE 2 framework [15], MGDrivE 3 incorporates modules for inheritance (the distribution of offspring genotypes for given maternal and paternal genotypes), mosquito life history (development from egg to larva to pupa to adult), landscape (the distribution and movement of mosquitoes through a metapopulation), and epidemiology (reciprocal pathogen transmission between mosquitoes and humans). MGDrivE 3 offers three substantial improvements beyond the functionality included in MGDrivE 2 - a sampling algorithm that allows decoupling of the mosquito and human model components, incorporation of a more detailed malaria transmission model, and inclusion of mosquito traps - each described in depth below. The software was developed using the R programming language, and retains the SPN formulation of the MGDrivE 2 package.

### Decoupled vector-host sampling framework

Decoupling the vector and host portions of the modeling framework is a major contribution of MGDrivE 3. Vector-borne disease models describe the reciprocal transmission of pathogens between vectors and hosts. Prior models have represented the vector and host state space as compartmental models represented by ordinary [16, 17, 44] or partial differential equations (PDEs) [10], or as individual-based models [19, 20]. In each of these models, vectors and hosts have the same state space and mathematical representation. In MGDrivE 3, the vector model is formulated as a SPN with a discrete state space, so we developed a sampling framework to allow the vector and host models to communicate, even if they have different representations.

Decoupling of vector-borne disease models is facilitated by the fact that all the vector model needs to know about the host model is the density and level of infectiousness in hosts, and vice versa. Communicating between the two model portions can therefore be accomplished by exchanging two composite parameters: i) the force of infection on hosts (*λ*_*H*_), i.e., the probability that a host is infected per unit time, and ii) the force of infection on vectors (*λ*_*V*_), i.e., the probability that a vector is infected per unit time. For malaria, *λ*_*H*_ is equal to the entomological inoculation rate (EIR, the number of infectious mosquitoes per human multiplied by their human biting rate) multiplied by the probability of the human becoming infected given an infectious bite. Similarly, *λ*_*V*_ is proportional to the human biting rate multiplied by the proportion of humans that are infectious multiplied by the probability of the mosquito becoming infected [21].

A schematic of an inter-model sampling algorithm for malaria, which we implement in this paper, is depicted in Fig 1. At each model iteration: i) the host model samples *λ*_*H*_ from the vector model, ii) the host model increments its infectious states for a time equal to one time step, iii) the vector model samples *λ*_*V*_ from the host model, and iv) the vector model increments its infectious states for one time step. While the MGDrivE 3 vector model is represented as an SPN with discrete state space, it is agnostic to how the host model is represented, which in this case is a system of PDEs with continuous state space [10, 11]. Additional considerations in implementing this algorithm include: i) choosing a time step appropriate to both models, ii) ensuring the MGDrivE 3 vector model produces output consistent with the vector model within the host model that it replaces, and iii) validating the EIR produced by the combined model framework.

**Fig 1.**
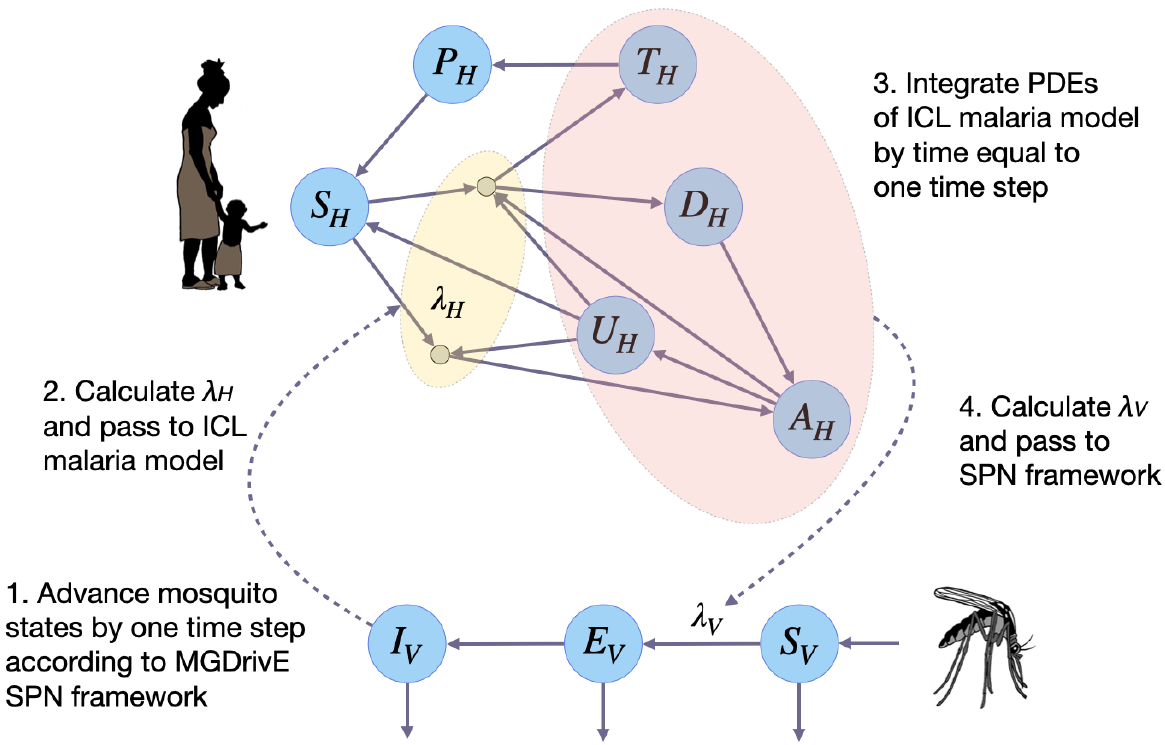
Schematic of decoupled vector-host sampling algorithm for malaria. MGDrivE 3 uses a stochastic Petri net framework to model progression of adult female mosquitoes from susceptible (*S*_*V*_) to exposed/latently infected (*E*_*V*_) to infectious for malaria (*I*_*V*_). This framework is linked to an adapted version of the Imperial College London (ICL) malaria model, which is represented as a set of partial differential equations. In the ICL model, humans progress from susceptible (*S*_*H*_) to either symptomatic or asymptomatic infection. Humans who develop a symptomatic infection and are either treated (*T*_*H*_) or diseased and untreated (*D*_*H*_). Treated humans advance to a prophylactic protection state (*P*_*H*_) and eventually become susceptible again. Untreated symptomatic humans develop successively lower-density infections, from symptomatic to asymptomatic but detectable by rapid diagnostic test (RDT) (*A*_*H*_) to asymptomatic and undetectable by RDT (*U*_*H*_). Asymptomatic humans can also be super-infected. To allow the two frameworks to communicate, at each time step: i) the ICL human model samples the force of infection in humans (*λ*_*H*_) from the MGDrivE 3 vector model, ii) the ICL human model increments its infectious states for a time equal to one time step, iii) the MGDrivE 3 vector model samples the force of infection in vectors (*λ*_*V*_) from the ICL human model, and iv) the MGDrivE 3 vector model increments its infectious states for one time step.

While a specific use case is presented in Fig 1, this inter-model sampling framework applies generally - it could equally be applied to models of arboviruses transmitted by *Aedes aegypti* [44], or to models of citrus greening disease transmitted by *Diaphorina citri* [22], provided the appropriate model adjustments are made.

### Malaria transmission model

Given the decoupled sampling algorithm, we incorporated an adapted version of the malaria model developed by the ICL malaria modeling group [10, 11] into MGDrivE 3. The MGDrivE 3 vector framework may be linked to several published malaria models; however, the ICL model represents a suitable level of parsimony for the current stage of development, as it can be described by a succinct set of PDEs while incorporating several important features of malaria epidemiology, and has been fitted to extensive malaria data sets throughout sub-Saharan Africa [10, 11]. Important epidemiological details captured in this model include symptomatic and asymptomatic infection, variable parasite density and superinfection in humans, human age structure, mosquito biting heterogeneity, and antimalarial drug therapy and prophylaxis. The model also includes several forms of immunity: i) pre-erythrocytic immunity reduces the probability of infection if bitten by an infectious mosquito; ii) acquired and maternal clinical immunity represent the effects of blood stage immunity on reducing the probability of developing clinical symptoms and severe illness; and iii) detection immunity represents the effects of blood stage immunity on reducing the detectability of an infection and onward transmission to mosquitoes. A full set of equations describing the ICL malaria model is provided in the S1 Text.

In incorporating certain genetic vector control tools - e.g., gene drive systems intended to spread disease-refractory genes into mosquito populations [2, 23] - an important addition to the epidemiological framework is transmission parameters that are mosquito genotype-specific. In the ICL malaria model, the force of infection on humans, *λ*_*H*_ (*a, t*), is dependent on both age, *a*, and time, *t*, and is a product of the EIR, *ε*(*a, t*), and the transmission probability, *b*, i.e.:

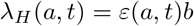

A given human could be bitten by a mosquito having any genotype, *g*, from the set of all genotypes, 𝒢, proportional to its time-varying frequency in the population, *p*_*g*_(*t*). For an effector gene that blocks mosquito-to-human transmission, the transmission probability, *b*_*g*_, will be genotype-specific, and so the expected transmission probability is equal to the time-varying weighted average:

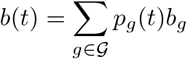

Incorporating more epidemiological detail into the model of transmission dynamics also allows more nuanced epidemiological outcomes to be calculated. As the ICL malaria model is age-stratified, both malaria prevalence and incidence can be calculated according to age group. Prevalence is calculated across all infectious human compartments - treated and untreated symptomatic disease, asymptomatic but detectable by rapid diagnostic test (RDT), and asymptomatic but undetectable by RDT - since each of these compartments contributes to onwards transmission of malaria to mosquitoes. Clinical incidence refers to new clinical malaria cases within a defined time interval, and is of particular relevance to the healthcare system. One commonly reported metric is malaria prevalence among children aged 2-10 years [24], as pediatric cases of malaria tend to be the most severe. A mathematical description of how each of these outcomes is calculated within the MGDrivE 3-ICL malaria model framework is provided in the S1 Text.

### Additional interventions and seasonality

Additional functionality has been included in both the vector and host portions of the MGDrivE 3 framework to incorporate currently-available interventions that genetic control tools would likely be implemented in conjunction with. While a range of novel vector control tools are currently under development [25]; the mainstay of malaria interventions for the last two decades has been a combination of LLINs, IRS and antimalarial drugs - largely artemisinin-combination therapy (ACT). Some combination of these interventions will invariably be present when a genetic control intervention is implemented, and it is important to characterize their implications for both vector population dynamics and vector-borne disease transmission. We model the impact of LLINs and IRS on mosquito life history parameters according to the elaborated feeding cycle model developed by Le Menach *et al*. [26] and adapted by Griffin *et al*. [10].

Within this framework, LLINs and IRS increase the mortality rate and decrease the biting rate of adult mosquitoes, and also decrease the egg-laying rate by virtue of extending the gonotrophic cycle. Equations for how each of these parameters are impacted by different coverage levels of LLINs and IRS are provided in the S1 Text. The proportion of symptomatic malaria cases that receive antimalarial drug therapy is included within the ICL malaria model [10, 11].

MGDrivE 3 also includes updated functionality for incorporating seasonal weather patterns. While MGDrivE 2 allows mosquito life history parameters, such as adult and larval development and mortality rates, to vary with time in response to environmental variables such as temperature and rainfall [15], the new framework utilizes environmental data to generate seasonal profiles to modulate these parameters. Rather than using raw daily rainfall data, which varies from year to year, the Umbrella R package [27] is used to fit a mixture of sinusoids to the rainfall data. This provides a more general characterization of the seasonal trends at a given location, and facilitates comparison across other locations with similar seasonal patterns. As with MGDrivE 2, development times are Erlang-distributed, and the model of White *et al*. [28] is used to modulate larval carrying capacity and hence density-dependent mortality in response to recent rainfall - a key driver for *Anopheles* population dynamics.

### Traps and spatial surveillance

In MGDrivE 3, the landscape module of MGDrivE 2 has been extended to accommodate traps as part of the mosquito metapopulation. In MGDrivE and MGDrivE 2, the landscape module describes the distribution of mosquitoes across discrete, randomly-mixing population nodes, with movement between them quantified by a dispersal kernel [5, 15]. MGDrivE 3 additionally accommodates “trap nodes” in one of two ways: i) traps are placed within a subset of population nodes, and are associated with a probability of trapping for mosquitoes within the corresponding population node per unit time, and ii) traps are assigned their own nodes, and are associated with coordinates and an attractiveness kernel, which includes an amplitude, mean distance of attractiveness, and other parameters as required by the kernel function. The former case is appropriate for applications on a larger spatial scale (e.g., where population nodes represent villages that traps may be placed in), while the latter is appropriate for applications on a finer spatial scale (e.g., where nodes represent blood, sugar or water sources that traps are placed relative to). In both cases, the landscape including traps may be generated using MGSurvE (Mosquito Gene Surveillance) [29]. Here, the number and locations of traps may be user-specified, along with their trapping probabilities (for the former case) or attractiveness kernel parameters (for the latter case), which should be chosen according to the types of traps being modeled.

Data analysis functions are provided to visualize the distribution of mosquitoes having certain genotypes that are captured by each trap over time.

## Results

To demonstrate how MGDrivE 3 can be used to simulate releases of gene drive-modified mosquitoes, including implications for epidemiological outcomes and surveillance, we have provided examples and information on GitHub at https://github.com/amondal2/MGDrivE3-Examples/tree/main/examples. In the first example, we model the release of a full gene drive system designed to drive malaria-refractory genes into an *An. coluzzii* mosquito population with seasonal population dynamics, pre-existing interventions and transmission intensity calibrated to a setting resembling the island of São Tomé, São Tomé and Príncipe. The full gene drive system resembles one engineered in *An. coluzzii* [2], which includes dual linked effector genes targeting the malaria pathogen, and is one of the most promising population replacement systems in a mosquito vector to date. While we model this system in a setting chosen largely for its isolation [30], we note that regulatory and biosafety issues must be considered seriously for self-propagating systems with the potential to spread beyond their release site [31].

Four alleles are considered at the gene drive locus: an intact drive allele containing disease-refractory genes (denoted by “H”), a wild-type allele (denoted by “W”), a functional, cost-free resistant allele (denoted by “R”), and a non-functional or otherwise costly resistant allele (denoted by “B”). The inheritance dynamics of this system were fitted to laboratory cage data and are provided in Carballar-Lejarazú *et al*. [2] with model parameters summarized in Table S1. Notably, we considered a 10% fitness cost associated with mosquitoes carrying either one or two copies of the intact drive allele, as there were no major fitness loads inferred in the *An. coluzzii* cage experiments [2]; however, fitness costs due to integration and expression of the gene drive system could become apparent in the field. Additionally, we assume that mosquitoes carrying either one or two copies of the H allele confer complete mosquito-to-human transmission blocking, consistent with data from Carballar-Lejarazú *et al*. [2] for sporozoite thresholds *≥* 7, 500.

The life history module is parameterized with typical bionomic parameters for *An. coluzzii* [28, 32], *with incorporation of a generalized seasonal profile that modulates certain life history parameters. In MGDrivE 3, as in MGDrivE 2, the carrying capacity of the environment for larvae is a function of recent rainfall, and a mathematical relationship from White et al*. [28] is used to translate local rainfall data to larval carrying capacity; however, in this example, we capture broad variations in the rainfall profile of São Tomé and Príncipe using the umbrella [27] package in R, using a shapefile of the national administrative boundary and a three-year timeframe for rainfall data (Fig 2). Otherwise, the life history module mirrors that of MGDrivE 2, including mean-variance relationships describing development times of the juvenile life stages [33]. For the purpose of this demonstration, and to emphasize the novel epidemiological component of MGDrivE 3, the island of São Tomé was treated as a single randomly mixing population.

**Fig 2.**
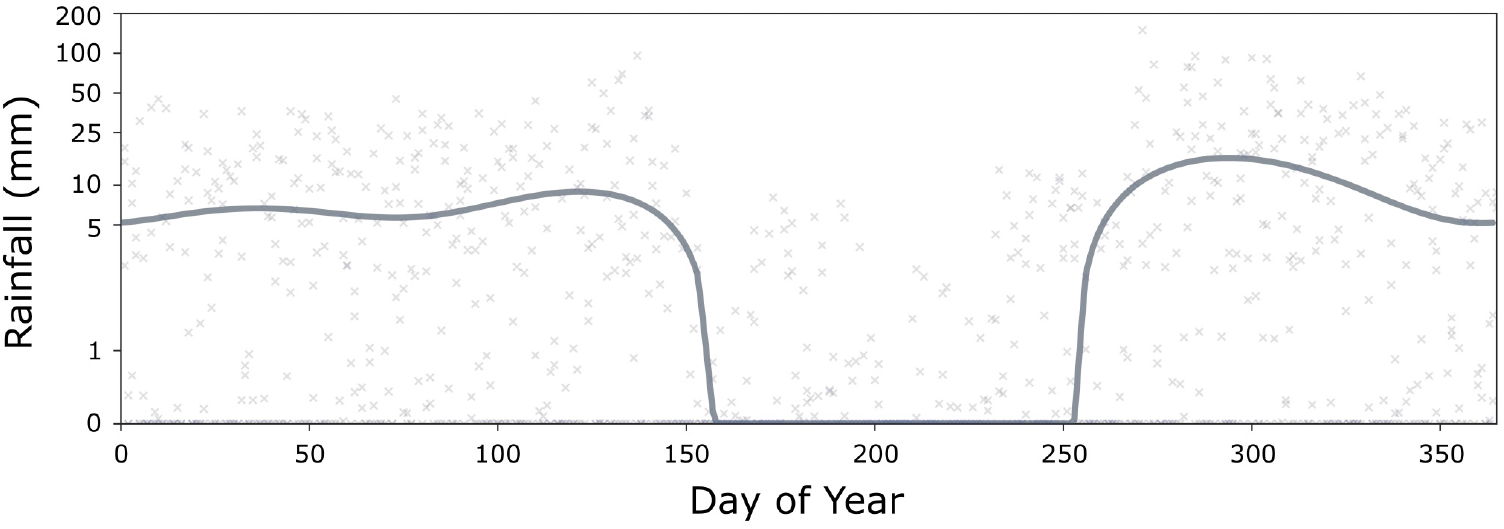
Seasonal rainfall profile for São Tomé and Príncipe. Points represent mean daily rainfall measurements (in mm) for the three years between January 1st, 2017 and Dececember 31st, 2019. The solid line represents the seasonal rainfall profile, fitted using the umbrella [27] package in R, used to calculate the time-varying environmental carry capacity for larvae in the life history module of MGDrivE 3.

The ICL malaria model is parameterized according to published intervention coverage and transmission levels for São Tomé and Príncipe - an LLIN coverage of 62%, IRS coverage of 66.5%, 2% of symptomatic malaria cases being treated with antimalarial drugs, and an all-ages *P. falciparum* prevalence of 2%, according to the World Health Organization Global Health Observatory (https://www.who.int/data/gho). The LLIN and IRS coverage parameters modify vector parameters in the life history module, while the antimalarial treatment parameter is input directly into the ICL model. Output from the ICL malaria model is then calibrated to all-ages malaria prevalence in the context of interventions and the seasonal rainfall profile by multiplying the carrying capacity for larvae by a constant. Other parameters of the ICL malaria model describe heterogeneity, human infectious periods, various types of immunity and treatment, and are as described in the original model [10, 11]. Finally, we note that these simulations are intended to demonstrate the software’s capabilities and that, while the simulations are calibrated to data from São Tomé, they are not intended to provide an accurate forecast of gene drive dynamics on the island, or to imply approval of the intervention by the local population and regulatory agencies.

### Simulation workflow

The code for running this simulation is available at: https://github.com/amondal2/MGDrivE3-Examples/blob/main/examples/stp local.R. We begin by loading the MGDrivE and MGDrivE 2 packages in R to gain access to the inheritance cubes, mosquito life history and malaria modeling functionality required for the simulation. Next, we load the inheritance cube for the TP13 population replacement gene drive system in *An. coluzzii* [2]. This specifies the inheritance-biasing properties of the system, as well as its malaria transmission-blocking effect. Note that there are a variety of other inheritance cubes available in the MGDrivE software - e.g., *Wolbachia* [34], release of insects carrying a dominant-lethal gene (RIDL) [35], precision-guided sterile insect technique (pgSIT) [3], population suppression gene drive [1], and remediation systems such as ERACR (element for reversal of the autocatalytic chain reaction) [36] - and users are also able to design their own inheritance cubes. Next, we specify general simulation parameters, such as the simulation length, the timestep of the stochastic model, and the timestep at which data is output (daily). Fitted reproductive fitness parameters for the TP13 construct in *An. coluzzii* [2] are loaded, and a 10% fitness load on male mating competitiveness and female fecundity is implemented, as described earlier.

Next, we specify details of the epidemiology module - baseline malaria prevalence, human population size, human age stratification, and coverage levels of LLINs, IRS and antimalarial drugs. Following this, as with MGDrivE 2, the “places” and “transitions” of the SPN formulation are set up using the “spn P epi decoupled node()” and “spn T epi decoupled node()” functions. Equilibrium values of states in the mosquito and human models are calculated using the “equilibrium Imperial decoupled()” function, and as the ICL malaria model requires the annual EIR to calculate the state distribution at equilibrium, a function is provided to convert malaria prevalence to EIR.

Next, the seasonal rainfall profile used to calculate the larval carrying capacity time-series (described above) is used to calculate time-varying hazard rates for density-independent larval mortality. Custom time-varying hazard functions for larval mortality are provided, and hazard functions are provided for the mosquito life history and ICL malaria transmission models. The MGDrivE 2 vignette, “Simulation of Time-inhomogeneous Stochastic Processes,” provides instructions for writing user-specified time-varying hazard functions. Finally, we specify the release scheme - genotypes, size and timing of releases - using an “events” dataframe.

With all model components specified, we call the “sim trajectory R decoupled()” function to simulate model trajectories. This implements a tau-leaping algorithm to sample stochastic trajectories, and records daily output to an R dataframe. For further analysis external to R, we provide functionality to write simulation output to CSV files.

### Entomological dynamics

In Fig 3, we display a potential visualization scheme for the entomological and epidemiological outcomes of the simulated gene drive mosquito release described above. This figure was generated in Python and is available at https://github.com/amondal2/MGDrivE3-Examples/tree/main/viz. We note that MGDrivE 3 is not dependent on Python, and the MGDrivE 3 R package provides basic plotting and analysis functions for model output visualization. In this case, we generated data for 15 stochastic model repetitions, and the dynamics displayed in Fig 3 depict the mean output of these repetitions. Fig 3A depicts allele frequencies for adult female mosquitoes over the simulation period. After eight consecutive releases of 20,000 male mosquitoes homozygous for the TP13 construct (H), the H allele rapidly spreads through the population, reaching near-fixation within a few months. This is a result of the high accurate homing rate, as determined by laboratory experiments [2], relatively low fitness costs (estimated), and low rate of resistance allele generation.

**Fig 3.**
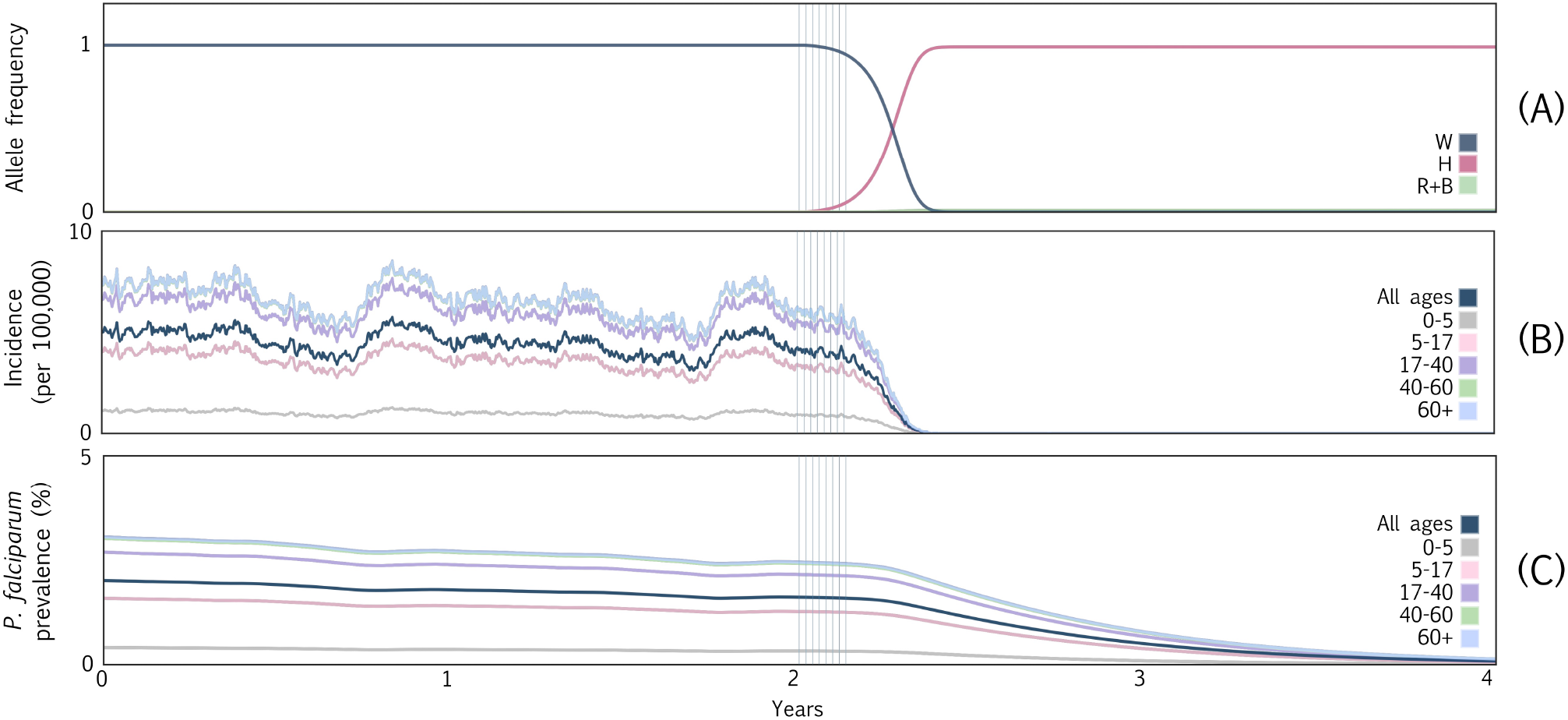
Example MGDrivE 3 simulations for a full gene drive system designed to drive dual malaria-refractory genes into an *An. coluzzii* mosquito population with seasonal population dynamics, transmission intensity and interventions calibrated to a setting resembling the island of São Tomé, São Tomé and Príncipe. The gene drive system resembles one recently engineered in *An. coluzzii* [2] in which all drive components - the Cas9, guide RNA and effector genes - are all present at the same locus. Four alleles are considered: an intact drive allele (denoted by “H”), a wild-type allele (denoted by “W”), a functional, cost-free resistant allele (denoted by “R”), and a non-functional or otherwise costly resistant allele (denoted by “B”). Model parameters describing the construct, mosquito bionomics and malaria transmission are summarized in Table S1. **(A)** Allele frequencies for adult female mosquitoes over the simulation period. Grey vertical bars beginning at year two denote eight consecutive weekly releases of 20,000 male mosquitoes homozygous for both the gene drive construct. The high efficiency of the drive system and low rate of resistance allele generation mean that almost no disease-competent *An. coluzzii* mosquitoes remain five months after the release. **(B)** Daily clinical malaria incidence per 100,000 people partitioned according to age group. Reductions in human incidence within five months of the release parallel spread of the drive construct in the mosquito population. **(C)** *P. falciparum* malaria prevalence partitioned according to age group. As humans recover from infection and few develop new infections, the *P. falciparum* parasite rate declines until it reaches near undetectable levels by year five.

Homing-susceptible wild-type alleles (W) are quickly eliminated, although a small number of in-frame and out-of-frame resistance alleles (R and B, respectively) accumulate since, although they are generated infrequently, they slightly outcompete the H alleles in terms of fitness. Note that while these dynamics represent a potential outcome of TP13 gene drive mosquito releases, the dynamics are highly dependent on the relative fitness of H and R/B allele-carrying mosquitoes, while are difficult to accurately quantify outside the field.

### Epidemiological dynamics

Here, we demonstrate the refined epidemiological outcomes obtained by linking the human portion of the ICL malaria transmission model to the vector portion of MGDrivE 3. We depict age-stratified clinical incidence in Fig 3B and age-stratified prevalence in Fig 3C. The rapid spread of the gene drive allele through the *An. coluzzii* population, and its strong modeled transmission-blocking effect, mean that humans are no longer exposed to new infectious mosquito bites five months after the beginning of the release schedule, and hence clinical incidence also falls to zero on this timescale.

Notably, clinical incidence includes symptomatic cases that are either treated or untreated (i.e., the *T*_*H*_ and *D*_*H*_ compartments in the ICL malaria model depicted in Fig 1), and does not include asymptomatic cases that are either detectable or undetectable by RDTs (i.e., the *A*_*H*_ and *U*_*H*_ compartments depicted in Fig 1). Stochastic variation in clinical incidence is pronounced due to the small number of incident cases relative to the total population.

São Tomé is a low-transmission setting with little acquired immunity, so incidence and prevalence are lower in younger age groups (0-5 and 5-17 years old) due to maternal immunity and the lesser skin surface area available for mosquito bites. *P. falciparum* prevalence includes all diseased states - i.e., symptomatic disease, whether treated or untreated (*T*_*H*_ and *D*_*H*_, respectively), and asymptomatic disease, whether detectable or undetectable by RDTs (*A*_*H*_ and *U*_*H*_, respectively). Prevalence in the human population takes much longer to decline than incidence, as an individual can harbor *P. falciparum* parasites for 1-2 years if left untreated [37], which is common for asymptomatic infections. These predictions highlight the transformative promise of gene drive interventions for malaria control; however, we caution that there are several limitations - notably, treatment of São Tomé as a panmictic population of humans and mosquitoes, calibration to malaria prevalence data that is likely underreported [38], and lack of knowledge of the fitness and transmission parameters of gene drive mosquitoes in the field, including their evolution over several years - which preclude the confidence with which such predictions can be made.

### Spatial surveillance

Finally, we demonstrate the capability of MGDrivE 3 to simulate surveillance of mosquitoes via traps placed throughout a landscape. The code for this example is available at https://github.com/amondal2/MGDrivE3-Examples/blob/main/examples/traps.R. We used the MGSurvE framework [29] to optimize the placement of five traps across a spatial network resembling the southern portion of São Tomé, São Tomé and Príncipe. This landscape is described by Sánchez C. *et al*. [29] - namely, nodes were sourced from the São Tomé and Príncipe census (https://projectsportal.afdb.org/dataportal/VProject/show/P-ST-KF0-001) and aligned with coordinates from Google Maps (https://www.google.com/maps). Daily mosquito movement probabilities were derived using an ecology-motivated algorithm [39], with model output calibrated to mark-release-recapture experiments on *An. gambiae* sensu lato [40, 41]. Traps were placed within population nodes to represent placement within selected villages, and trapping probabilities were specified, along with the rest of the landscape, in MGSurvE.

We consider a release in the south of the island and monitor the progression of gene drive phenotypes for trapped mosquitoes over time. As for the epidemiological simulation, we consider eight weekly releases of male *An. coluzzii* homozygous for the gene drive system. We consider a simplified version of the TP13 gene drive construct [2] with only a single resistance (R) allele. The cutting frequency at the target site for this construct is 1.0, and the rate of accurate homology-directed repair is 0.99. The inheritance cube is flexible to specify genotype-specific mating fitness, multipliers on adult mortality, male and female pupatory success, and reductions in fertility, but we do not modify them in this example. We model mosquitoes as accumulating in traps over the course of a week, after which they are counted and the traps are “reset.” We also tally gene drive phenotypes when trapped mosquitoes are counted, considering a marker allele associated with both the intact drive allele (H) and the wild-type target allele (W) [2]. This allows us to distinguish the following genotypes: HH/HR, WW/WR, HW, and RR. Fig 4 depicts the time-series of gene drive marker phenotypes in each trap by week, with the time of first detection of a transgenic mosquito indicated by a vertical line for each trap. Output like this will be useful to model surveillance strategies for the progression of field trials and interventions, and the emergence of alternative alleles that could interfere with intervention effectiveness [14].

**Fig 4.**
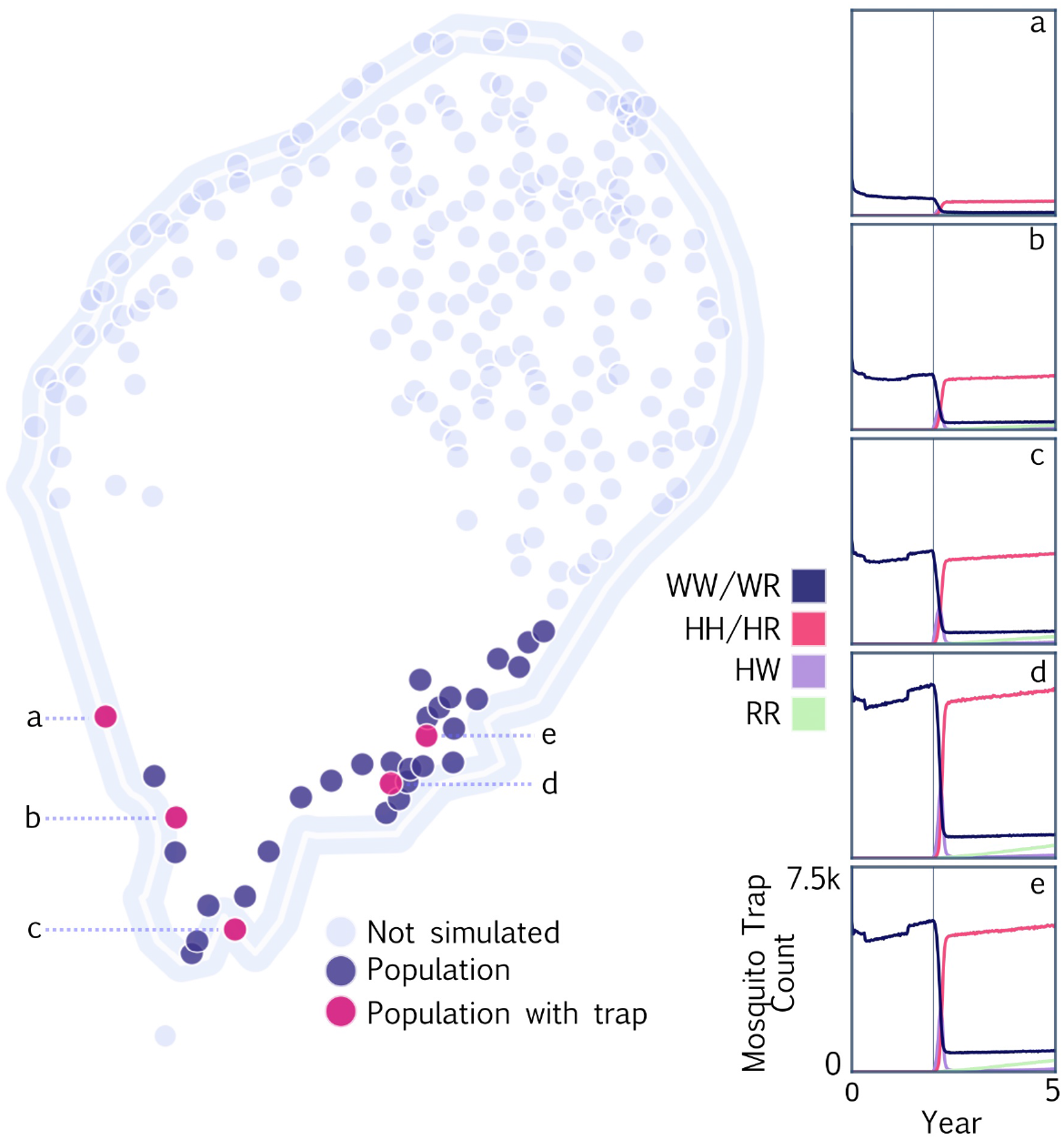
Example MGDrivE 3 simulations for spatial surveillance of a full gene drive system on the island of São Tomé, São Tomé and Príncipe. Mosquito population nodes represent villages and suburbs of comparable size with mosquito movement probabilities between localities derived from an ecology-motivated algorithm [39] and calibrated to mark-release-recapture data [40, 41]. Simulation was restricted to the southern portion of the island, with population nodes including traps depicted in pink and other population nodes depicted in blue. Traps were placed using the MGSurvE framework [29]. Eight weekly releases of a full gene drive system (cutting rate of 1.0 and homology-directed repair rate of 0.99) were simulated in the south of the island, and the phenotype distribution of trapped mosquitoes is depicted for the five trap nodes in panels a-e. Vertical lines denote the time of first transgene detection for each trap.

## Availability and future directions

MGDrivE 3 is available at https://cran.r-project.org/package=MGDrivE2 as version 2.1.0 of the MGDrivE 2 package, due to naming conventions. The source code is under the GPL3 License and is free for other groups to modify and extend as needed. Mathematical details of the model formulation are available in the S1 Text. Examples for running MGDrivE 3 simulations are available at https://github.com/amondal2/MGDrivE3-Examples/tree/main/examples, and documentation for MGDrivE 3 functions are available at the MGDrivE 2 project website at https://marshalllab.github.io/MGDrivE/docs_v2/index.html. To run the software, we recommend using R version 3.1.0 or higher.

We are continuing development of the MGDrivE 3 software package and welcome suggestions and requests from the research community regarding future directions. As gene drive mosquito projects advance from the lab to the field, we intend our software to address the evolving modeling needs of the technology [42] - from contributing to TPPs [4] and environmental risk assessments [43], to planning field trials, interventions [6, 7] and surveillance programs [14]. The epidemiological extensions offered in MGDrivE 3 will enable more accurate predictions of implications of mosquito genetic control for disease transmission, which are relevant as an outcome for TPPs, and field trial and intervention planning. This will also enable prediction of the impact of genetic control interventions alongside other currently-implemented interventions such as LLINs, IRS and ACTs. The surveillance extensions included in MGDrivE 3 will enable assessment of mosquito trapping schemes to both: i) measure the effectiveness of genetic control strategies in the field, and ii) detect unintended spread of gene drive alleles beyond field sites, and the emergence of alternative alleles broadly [14].

Logistical modeling questions are invariably associated with larger state spaces - more genotypes to keep track of, more human and mosquito disease states, and larger metapopulation networks - which quickly approach the computational limits of the modeling framework. To address this, we are exploring numerical sampling algorithms to increase computational efficiency and speed, and the use of lower-level programming languages such as C++. We are also interested in linking the vector portion of MGDrivE 3 to other epidemiological models that capture human transmission dynamics more comprehensively - e.g., dengue models that incorporate multiple serotypes with temporary cross-protective immunity and complications related to antibody-dependent enhancement [44], and individual-based malaria transmission models that allow sources of heterogeneity to be incorporated more comprehensively and for infection history to be directly associated with immune status [45]. There are also opportunities to adapt the framework to species of relevance to agriculture and conservation - e.g., enhanced epidemiological capabilities could be applied to citrus greening disease transmitted by *D. citri* [22], and surveillance functionality could be suitable for models of invasive rodents on islands [46].

## Acknowledgments

We thank Drs. Sean Wu, Rodrigo Corder, and Peter Winskill for comments on the manuscript and model development. We thank Prof. Azra Ghani and the malaria modeling group at Imperial College London for permission to link an adapted version of their malaria transmission model to our mosquito genetic control model.

## Data availability

MGDrivE 3 is available at https://cran.r-project.org/package=MGDrivE2 as version of the MGDrivE 2 package. The source code is under the GPL3 License and is free for other groups to modify and extend as needed. Examples for running MGDrivE 3 simulations are available at https://github.com/amondal2/MGDrivE3-Examples/tree/main/examples, and documentation for MGDrivE 3 functions are available at https://marshalllab.github.io/MGDrivE/docsv2/index.html.

## Supporting information

**Table S1. Model parameters describing the gene drive construct, mosquito bionomics and malaria epidemiology for simulations resembling releases on São Tomé, São Tomé and Príncipe**.

**S1 Text. Description of the modeling framework**. A description of the mathematical equations that govern the malaria transmission model, including prior interventions and outcomes of interest.

