## Supplementary material for "MGDrivE 3: A decoupled vector-human framework for epidemiological simulation of mosquito genetic control tools and their surveillance": S1 Text

#### S1 Text: Description of the modeling framework

Agastya Mondal[1,*], Héctor M. Sánchez C.[1], John M. Marshall [1,*]

**1** Divisions of Epidemiology and Biostatistics, School of Public Health, University of California, Berkeley, CA 94720, USA.

*

### Malaria transmission model

Malaria transmission is a complex process involving dynamics between host, parasite and vector, with influence from the environment. While the dynamics of MGDrivE [1] and MGDrivE 2 [2] focus on vector dynamics, several models are available that focus on pathogen transmission in the host [3, 4]. In MGDrivE 3, we incorporate an adapted version of the Imperial College London (ICL) malaria transmission model [4], as it represents a suitable level of parsimony and has been fitted to extensive malaria data sets throughout sub-Saharan Africa. The ICL malaria model contains several important components:

1. Time and age-structured equations describing the movement of humans into various disease states;
2. Equilibrium distribution based on baseline entomological innoculation rate (EIR) and age structure of the population; and
3. Population-level immunity functions which modulate various infection probabilities.

The state space is modeled as a set of partial differential equations (PDEs). The infection states are: susceptible (S), treated symptomatic disease (T), untreated symptomatic disease (D), asymptomatic infection that is detectable by rapid diagnostic test (RDT) (A), sub-patent infection that is undetectable by RDT (U), and post-treatment prophylaxis (P). The force of infection on humans (which depends on the EIR) is denoted $\Lambda$, the probability that symptoms develop after an infectious challenge is denoted $\Phi$, and the fraction of clinical cases that receive effective treatment is denoted $f_T$. The set of human state PDEs is shown below, with $a$ representing age and $t$ representing time.

$$\frac{\partial S}{\partial t} + \frac{\partial S}{\partial a} = -\Lambda S + \frac{P}{d_P} + \frac{U}{d_U}$$
$$\frac{\partial T}{\partial t} + \frac{\partial T}{\partial a} = \Phi f_T \Lambda (S + A + U) - \frac{T}{d_T}$$
$$\frac{\partial D}{\partial t} + \frac{\partial D}{\partial a} = \Phi(1 - f_T)\Lambda (S + A + U) - \frac{D}{d_D}$$
$$\frac{\partial A}{\partial t} + \frac{\partial A}{\partial a} = (1 - \Phi)\Lambda (S + U) + \frac{D}{d_D} - \Phi\Lambda A - \frac{A}{d_A}$$
$$\frac{\partial U}{\partial t} + \frac{\partial U}{\partial a} = \frac{A}{d_A} - \frac{U}{d_U} - \Lambda U$$
$$\frac{\partial P}{\partial t} + \frac{\partial P}{\partial a} = \frac{T}{d_T} - \frac{P}{d_P}$$

Here, $d_i$ indicates the mean duration of state $i$. Additionally, the model includes four forms of population-level immunity:

- Pre-erythrocytic immunity, $I_B$, reduces the probability of infection if bitten by an infectious mosquito;
- Acquired and maternal clinical immunity, $I_{CA}$ and $I_{CM}$, represent the effects of blood stage immunity in reducing the probability of developing clinical symptoms; and
- Detection immunity, $I_D$, represents the effect of blood stage immunity in reducing the detectability of an infection and onward transmission to mosquitoes.

The PDEs describing immunity are below. Note that $\varepsilon$ represents the EIR, $u_i$ limits the rate at which immunity can be boosted at high exposure for immunity state $i$, and $d_i$ determines the duration of immunity for immunity state $i$.

$$\frac{\partial I_B}{\partial t} + \frac{\partial I_B}{\partial a} = \frac{\varepsilon}{\varepsilon u_B + 1} - \frac{I_B}{d_B}$$
$$\frac{\partial I_{CA}}{\partial t} + \frac{\partial I_{CA}}{\partial a} = \frac{\Lambda}{\Lambda u_c + 1} - \frac{I_{CA}}{d_{CA}}$$
$$\frac{\partial I_{CM}}{\partial t} + \frac{\partial I_{CM}}{\partial a} = \frac{-I_{CM}}{d_{CM}}$$
$$\frac{\partial I_D}{\partial t} + \frac{\partial I_D}{\partial a} = \frac{\Lambda}{\Lambda u_d + 1} - \frac{I_D}{d_D}$$

Each immunity function is transformed to a reduction in the appropriate infection probability via a Hill function.

Instead of numerically solving the PDEs directly, we first discretize the model by age category and biting heterogeneity. To discretize by age, we augment each infection state by an age category. For example, if we had two age categories 0-10 years and 10-100 years, then we would have susceptible compartments $S_1$ and $S_2$, where $S_1$ contains the people in the 0-10 year category and $S_2$ contains the people in the 10-100 year category. This would apply for all infection states. In addition, each compartment contains a rate at which people age and therefore move between age compartments.

Then, each PDE becomes a discrete ODE representing an age compartment. For example,

$$\frac{dS_i}{dt} = -\Lambda S_i + \frac{P_i}{d_P} + \frac{U_i}{d_U} - \eta_i S_i + \eta_{i-1} S_{i-1}$$

gives the rate equation for the susceptible (S) state for age category $i$ where $\eta_i$ gives the aging rate from $S_i \longrightarrow S_{i+1}$ and similarly $\eta_{i-1}$ gives the aging rate from $S_{i-1} \longrightarrow S_i$. For the youngest age group, the $\eta_{i-1}S_{i-1}$ term would be left out, and for the oldest age group, the $\eta_i S_i$ term would be left out.

One implementation note is that the model assumes a fixed latent period of 12 days after an infectious challenge from a mosquito, after which either symptoms develop or an asymptomatic infection proceeds. Because of this fixed delay, the equations are technically formulated as "delay differential equations," where the current state depends on the previous state.

To initialize the distribution of disease and immunity states, the model takes as input the baseline EIR, the age structure of the population, the proportion of treated cases, and baseline entomological parameters. Some of the mosquito life cycle parameters will vary in the presence of interventions, which will be described in the next section.

Finally, an important novel contribution of this work incorporates a model for genotype-specific transmission probabilities. In the gene drive context, it is important to understand how mosquitoes modified with a certain allele can affect disease transmission. In the traditional context [5], force of infection on humans ($\lambda_H$) is proportional to the EIR ($\varepsilon$) and the probability of successful infection upon biting ($b$). In the ICL malaria model, the force of infection is expanded to include a term corresponding to pre-erythrocytic immunity ($I_B$):

$$\lambda_H \propto \varepsilon b I_B$$

In our adapted model, we allow for varying transmission probabilities depending on the genotype distribution of circulating mosquitoes in the model. For example, we may consider a gene drive system in which the homozygous transgenic mosquito (denoted "HH") confers perfect infection blocking such that $b_{HH} = 0$, and where wildtype mosquitoes (denoted "WW") have an infection probability of 0.55 ($b_{WW} = 0.55$). Then, to calculate the total infection probability for any time point, $t$, we take the weighted average over the circulating proportion of infectious mosquitoes of each genotype ($p_{HH}$ and $p_{WW}$) at time $t$, i.e.:

$$b(t) = p_{WW}(t) b_{WW} + p_{HH}(t) b_{HH}$$

More generally, for a genotype set $\mathcal{G}$, we have the total infection probability as the weighted average:

$$b(t) = \sum_{g \in \mathcal{G}} p_g(t) b_g$$

where $p_g(t)$ represents the population frequency of mosquitoes having genotype $g$ at time $t$, and $b_g$ represents the infection probability for genotype $g$. With all of the above components in place, the epidemiology model is fully specified.

### Epidemiological outcomes

In this modeling framework, it is important to specify the epidemiological outcomes of interest. Generally, we are interested in clinical incidence of disease, which refers to the number of new symptomatic cases per day, and *P. falciparum pathogen prevalence* (*PfPr*), which refers to the proportion of the population harboring the malaria pathogen, whether symptomatic or asymptomatic. Because our model is age-structured, we can consider these outcomes for different age categories. Malaria outcomes are often more severe for younger individuals [6], so it makes sense to consider incidence and prevalence for younger age groups (e.g., 0-2 years) in order to better understand how gene drive interventions will mitigate these cases. Additionally, we can consider all-ages prevalence and incidence to understand how the intervention will perform in the entire population.

Mathematically, we define *PfPr* as the sum of all individuals in infectious disease states: symptomatic and treated (T), symptomatic and untreated (D), asymptomatic patent infection (A), and asymptomatic subpatent infection (U). Therefore, the all-ages (often denoted by the subscript 0-99 to denote the entire lifespan in years) pathogen prevalence at a given time point, $t$, is given by:

$$PfPr_{[0-99]}(t) = \sum_{a \in \mathcal{A}} (A_a(t) + U_a(t) + D_a(t) + T_a(t))$$

where $\mathcal{A}$ is the set of all age compartments. Similarly, the 0-2 years *PfPr* is given by:

$$PfPr_{[0-2]}(t) = A_{[0-2]}(t) + U_{[0-2]}(t) + D_{[0-2]}(t) + T_{[0-2]}(t)$$

As for clinical incidence, we first define some parameters:

- $\phi$: the probability of acquiring clinical disease upon infection (proportional to immunity levels via a Hill function);
- $\lambda_H$: the force of infection on humans (linearly proportional to the EIR, $\varepsilon$); and
- $Y$: the sum of non-clinical disease states, susceptible (S), asymptomatic patent infection (A), and subpatent infection (U).

Then we can define the all-ages clinical incidence as:

$$CI_{[0-99]}(t) = \sum_{a \in \mathcal{A}} \phi_a(t) \lambda_{H,a}(t) Y_a(t)$$

and the 0-2 years clinical incidence as:

$$CI_{[0-2]}(t) = \phi_{[0-2]}(t) \lambda_{H,[0-2]}(t) Y_{[0-2]}(t)$$

Generally, we are interested in these outcomes with respect to their baseline or pre-intervention values. In our analyses, we will calculate the reduction in prevalence and clinical incidence as our outcomes of interest. As we will be running many stochastic repetitions of the simulation for a given parameter set, the mean reduction over the repetition set and simulation timespan will be used. Note that in this formulation, each disease state is a proportion, with all disease states summing to 1. If instead we wish to model a population of $N_H$ humans, then we would simply divide each outcome by $N_H$ to obtain the proportional value.

#### Additional interventions

Here we show the full derivation of how indoor residual spraying (IRS), long-lasting insecticide-treated nets (LLIN), and artemisisin-based combination therapy (ACT) interventions modify baseline mosquito life cycle parameters. This derivation is adapted from previous work [4, 7].

First, we assume that, at baseline, we have three proportions of active vector control interventions, $\{\chi_{IRS}, \chi_{LLIN}, \chi_{ACT}\}$, which represent the proportion of humans in the model covered by the given intervention. Then, $\chi_{ACT}$ corresponds to the proportion of symptomatically infected humans that are treated upon infection, $f_T$.

Then, $\{\chi_{IRS}, \chi_{LLIN}\}$ jointly modify various mosquito life cycle parameters. First, we model the impact of LLINs and IRS on the length of the mosquito gonotrophic cycle (i.e., the time taken for a mosquito to take a blood meal and lay eggs before seeking its next blood meal). This time can be divided into $\tau_1$ (the time spent foraging) and $\tau_2$ (the time spent ovipositing and resting). Then, the length of the gonotrophic cycle in the presence of vector control is given by:

$$\frac{1}{\delta_c} = \frac{\tau_1(0,0)}{1-z} + \tau_2$$

where $\tau_1(0,0)$ represents the time time spent foraging with LLIN and IRS coverages of zero, and:

$$\begin{aligned} z = Q_0 c_{LLIN}\theta_B r_{LLIN} + Q_0 c_{IRS}\theta_I r_{IRS} + \\ Q_0 c_{LLIN,IRS}(\theta_I - \theta_B) r_{IRS} + \\ Q_0 c_{LLIN,IRS}\theta_B r_{IRS,LLIN} \end{aligned}$$

Here, $Q_0$ represents the human blood index, $\theta_B$ represents the proportion of bites taken on a person in bed, $\theta_I$ represents the proportion of bites taken on a person outdoors, $r_{IRS}$ represents the probability of repeating a feeding attempt in the presence of IRS, $r_{IRS,LLIN}$ represents the probability of repeating a feeding attempt in the presence of IRS and LLINs, and:

$$\begin{aligned} c_{LLIN} &= \chi_{LLIN} - \chi_{LLIN}\chi_{IRS} \\ c_{IRS} &= \chi_{IRS} - \chi_{LLIN}\chi_{IRS} \\ c_{LLIN,IRS} &= \chi_{LLIN}\chi_{IRS} \\ c_0 &= 1 - \chi_{LLIN} - \chi_{IRS} + \chi_{LLIN}\chi_{IRS} \end{aligned}$$

Then, with the modified gonotrophic cycle calculated ($\delta_C$), we can model the impact of LLINs and IRS on the adult mosquito death rate. We express the mortality rate in the presence of vector control as:

$$\mu_{V,C} = -\log p(\chi_{IRS}, \chi_{LLIN})$$

where $p$ represents the probability of an adult mosquito surviving one day. Then we can break down $p$ into two components $p_1$ (the probability of surviving the mosquito stage) and $p_2$ (the probability of surviving the blood meal stage):

$$p(\chi_{IRS}, \chi_{LLIN}) = (p_1(\chi_{IRS}, \chi_{LLIN})p_2)^{\delta_c}$$

where:

$$p_1(\chi_{IRS}, \chi_{LLIN}) = \frac{p_1(0,0)w}{1 - zp_1(0,0)}$$

$z$ is the same as above and $w$ gives the probability that a mosquito successfully feeds and survives a single feeding attempt:

$$\begin{aligned} w = 1 - Q_0 + Q_0 c_0 + Q_0 c_{LLIN}(1 - \theta_B + \theta_B s_{LLIN}) + \\ Q_0 c_{IRS}(1 - \theta_I + \theta_I s_{IRS}) + \\ Q_0 c_{IRS,LLIN}((\theta_B - \theta_I)s_{IRS} + 1 - \theta_I + \theta_B s_{LLIN,IRS}) \end{aligned}$$

Here, $s_{LLIN}$ and $s_{IRS}$ represent the probability of feeding and surviving in the presence of LLINs and IRS, respectively. The non-intervention survival probabilities are given by:

$$p_1(0,0) = e^{-\mu_V \tau_1(0,0)}$$
$$p_2 = e^{-\mu_V \tau_2}$$

Now, we have mathematical expressions for the gonotrophic cycle length and adult mortality rate ($\delta_c$ and $\mu_{V,c}$ respectively). We can finally model the impact of LLINs and IRS on the egg laying rate of the adult female mosquito. In the absence of vector control, the egg laying rate is given by:

$$\beta = \frac{\varepsilon \mu_V}{e^{\frac{\mu_V}{\delta}} - 1}$$

where $\varepsilon$ is the number of viable eggs laid per oviposition cycle. Then, with the previously defined parameters, the egg laying rate in the presence of vector control interventions is simply:

$$\beta_c = \frac{\varepsilon \mu_{V,c}}{e^{\frac{\mu_{V,c}}{\delta_c}} - 1}$$

Finally, we can modify the human biting rate per mosquito in the presence of LLINs and IRS. In the absence of interventions, the biting rate is given by:

$$a_V = \delta Q_0$$

The biting index under intervention is given by:

$$Q_c = 1 - \frac{1 - Q_0}{w}$$

where $w$ is the calculated probability from above. Then, using the modified gonotrophic cycle length previously derived ($\delta_c$), the modified biting rate is thus:

$$a_{V,c} = \delta_c Q_c$$

With these definitions in place, we have fully specified the impact of vector control interventions on mosquito life cycle parameters.
