## Supplementary material for "MGDrivE 3: A decoupled vector-human framework for epidemiological simulation of mosquito genetic control tools and their surveillance": Table S1

**Table S1.** Model parameters describing the gene drive construct, mosquito bionomics and malaria epidemiology for simulations resembling releases on São Tomé, São Tomé and Príncipe.

| Symbol: | Parameter: | Value: | Reference: |
|---|---|---|---|
| **Gene drive construct:** | | | |
| $f_{HH}$ | Homozygous fitness (relative to wildtype) on female mosquitoes | 0.9 | [1] |
| $f_H$ | Hemizygous fitness (relative to wildtype) on female mosquitoes | 0.9 | [1] |
| $m_{HH}$ | Homozygous male mating competitiveness | 1.05 | [1] |
| $m_H$ | Hemizygous male mating competitiveness | 1.78 | [1] |
| $p^M{}_W$ | Probability of wildtype allele staying intact across one generation in male mosquitoes | 0.0212 | [1] |
| $p^M{}_H$ | Probability of wildtype allele converting to H allele across one generation in male mosquitoes | 0.979 | [1] |
| $p^F{}_W$ | Probability of wildtype allele staying intact across one generation in female mosquitoes | 0.0015 | [1] |
| $p^F{}_H$ | Probability of wildtype allele converting to H allele across one generation in female mosquitoes | 0.985 | [1] |
| $p^{HH}{}_W$ | Probability of wildtype allele staying intact across one generation in gravid, homozygous female mosquitoes | 0.938 | [1] |
| $p^{HH}{}_R$ | Probability of wildtype allele converting to R allele across one generation in gravid, homozygous female mosquitoes | 0.0122 | [1] |
| $p^{HH}{}_B$ | Probability of wildtype allele converting to B allele across one generation in gravid, homozygous female mosquitoes | 0.0437 | [1] |
| $p^H{}_W$ | Probability of wildtype allele staying intact across one generation in gravid, hemizygous female mosquitoes | 0.997 | [1] |
| $p^H{}_R$ | Probability of wildtype allele converting to R allele across one generation in gravid, hemizygous female mosquitoes | 0.0007 | [1] |
| $p^H{}_B$ | Probability of wildtype allele converting to B allele across one generation in gravid, hemizygous female mosquitoes | 0.0017 | [1] |
| $b_{WW}$ | Wildtype mosquito-to-human transmission probability | 0.55 | [1] |
| $b_H$ | TP13 drive mosquito-to-human transmission probability | 0 | [1] |
| $c$ | Human-to-mosquito transmission probability | 0.15 | [1] |
| **Vector biology:** | | | |
| $\beta$ | Egg production per adult female (per day) | 21 | [2] |
| $T_E$ | Mean duration of egg stage (days) | 3 | [2] |
| $T_L$ | Mean duration of larval stage (days) | 7 | [2] |
| $T_P$ | Mean duration of pupal stage (days) | 1 | [2] |

| | | | |
|---|---|---|---|
| $CV(T_E)$ | Coefficient of variation, egg stage | 0.2 | [3] |
| $CV(T_L)$ | Coefficient of variation, larval stage | 0.3 | [3] |
| $CV(T_P)$ | Coefficient of variation, pupal stage | 0.2 | [3] |
| $K$ | Larval carrying capacity | Time-varying | [4] |
| $\mu$ | Adult mosquito mortality rate | Time-varying | [4] |
| $f$ | Blood feeding rate | 1/3 | [5] |
| $Q$ | Human blood index | 0.9 | [5] |
| **Vector control:** | | | |
| $\theta_B$ | Proportion of bites on a person in bed | 0.89 | [6] |
| $\theta_I$ | Proportion of bites on a person outdoors | 0.97 | [6] |
| $r_{LLIN}$ | Probability of repeating a feeding attempt in the presence of long-lasting insecticide-treated nets | 0.56 | [6] |
| $r_{IRS}$ | Probability of repeating a feeding attempt in the presence of indoor residual spraying | 0.60 | [6] |
| $s_{LLIN}$ | Probability of feeding and surviving in the presence of long-lasting insecticide-treated nets | 0.03 | [6] |
| $s_{IRS}$ | Probability of feeding and surviving in the presence of indoor residual spraying | 0 | [6] |
| **Intervention setting and demography:** | | | |
| $N_H$ | Human population size | 223,000 | [7] |
| $PfPr$ | All-ages *P. falciparum* prevalence | 0.02 | [8] |
| $\chi_{LLIN}$ | Proportion of population using long-lasting insecticide-treated nets | 0.62 | [7] |
| $\chi_{IRS}$ | Proportion of population using indoor residual spraying | 0.665 | [7] |
| $f_T$ | Proportion of population using artemisinin-based combination therapy | 0.02 | [7] |
